# Stress-induced transcriptional readthrough into neighboring genes is linked to intron retention

**DOI:** 10.1101/2022.03.24.485601

**Authors:** Shani Hadar, Anatoly Meller, Reut Shalgi

## Abstract

Exposure to certain stresses leads to readthrough transcription. Here we found that readthrough transcription often proceeds into the proximal gene downstream, in a phenomenon termed “read-in”. Using polyA-selected RNA-seq data from mouse fibroblasts, we identified widespread read-in in heat shock, oxidative and osmotic stress conditions. We found that read-in genes share distinctive genomic characteristics; they are extremely short, and highly GC rich. Furthermore, using ribosome footprint profiling we found that translation of read-in genes is significantly reduced. Strikingly, read-in genes show extremely high levels of intron retention during stress, mostly in their first intron, which is not explained by features usually associated with intron retention, such as short introns and high GC content. Finally, we found that first introns in read-in genes have weaker splice sites. Our data portray a relationship between read-in and intron retention, suggesting it may have co-evolved to facilitate reduced translation of read-in genes during stress.

## Introduction

Gene expression is known to be highly regulated in response to stress(Lopez-Maury et al., 2008). In the past decade, it is becoming increasingly clear that post-transcriptional RNA processing is also tightly regulated in response to stress (Biamonti and Caceres, 2009), including splicing (Nevo et al., 2012; Sabath et al., 2020; Shalgi et al., 2014), and polyadenylation (Di Giammartino et al., 2013). In particular, intron retention is dynamically regulated in various environmental and physiological conditions (Boutz et al., 2015; Shalgi et al., 2014; Wong et al., 2013). In heat shock, for example, widespread intron retention in more than a thousand genes was shown to occur, leading to the accumulation of polyadenylated, stable, intron-containing mRNAs in the nucleus (Shalgi et al., 2014). Indeed, one of the prevalent fates of intron-containing mRNAs is the prevention of nuclear export, leading to nuclear retention (Monteuuis et al., 2019).

More recently, it was found that several stress conditions, including heat shock, osmotic, oxidative stress (Vilborg et al., 2015; Vilborg et al., 2017), and hypoxia (Wiesel et al., 2018), lead to pervasive transcriptional readthrough, resulting in long continuous transcripts that can extend up to thousands of kilobases downstream to gene ends, and affecting thousands of genes in human and mouse cells (Vilborg et al., 2015; Vilborg et al., 2017). This phenomenon, which also happens during viral infection (Rutkowski et al., 2015; Zhao et al., 2018), is thought to occur due to reduced efficiency of polyadenylation(Rosa-Mercado and Steitz, 2021). Nevertheless, although it was shown to be regulated, and not due to a random failure (Vilborg et al., 2017), and even though several characteristics of readthrough genes have been identified (Vilborg et al., 2017), the underlying selectivity of stress-induced transcriptional readthrough, still remains elusive. Key genomic characteristics of readthrough-affected genes include depletion of polyadenylation motifs downstream to gene ends (Rutkowski et al., 2015; Vilborg et al., 2015; Vilborg et al., 2017), open chromatin marks past gene ends (Hennig et al., 2018; Vilborg et al., 2017), and close proximity to neighboring genes (Vilborg et al., 2017).

While several underlying pathways have been found to contribute to transcriptional readthrough during viral infection (Wang et al., 2020), and in osmotic stress (Rosa-Mercado et al., 2021), the consequences of this fairly newly identified phenomenon are still largely a mystery. Antisense regulation by readthrough transcripts has been proposed as one potential consequence (Vilborg et al., 2017), and was demonstrated to occur during senescence (Muniz et al., 2017). Interestingly, using nascent RNA-seq performed during HSV-1 viral infection, it was observed that readthrough transcription can extend into neighboring genes, which were therefore termed read-in genes (Rutkowski et al., 2015). Read-in was also shown to occur during Influenza virus infection (Roth et al., 2020). Nevertheless, whether these chimeric read-in transcripts can be detected as mature mRNAs, and what their potential fate is, has remained unknown.

Here we show that stress induces widespread readthrough into neighboring read-in genes, which is evident by the presence of mature, polyadenylated, mRNA transcripts in RNA-seq data. Read-in genes tend to reside close to their upstream readthrough genes, as expected. However, read-in is not simply a function of distance from the readthrough gene, as there are genes proximal to readthrough genes that do not show read-in. We found that read-in genes have distinct genomic characteristics; they are substantially short, they have less introns and their introns are shorter, and they tend to be GC rich. Additionally, genes with a high degree of read-in transcription are largely translationally inhibited. Importantly, we found that read-in genes display marked intron retention. Our analyses showed that although introns of read-in genes share characteristics associated with intron retention, namely they are short and GC rich, the high degree of intron retention observed could not be simply explained by these properties. As intron retention is known to be associated with nuclear retention, we speculate that these properties were evolutionarily retained, and therefore facilitate nuclear retention of read-in transcripts, thereby preventing potential aberrant translation resulting from readthrough of chimeric read-in genes during stress conditions.

## Results

### Stress induces transcriptional readthrough into neighboring genes

To compare the transcriptional and translational responses to stress, we analyzed polyA-selected mRNA-seq, as well as ribosome footprint profiling (Ribo-seq), performed in mouse NIH3T3 fibroblasts subjected to heat shock (42 degrees), oxidative stress (H2O2), or osmotic stress (KCl), for acute (2h) or sustained (7-8h) treatments. Comparison of the fold changes at the level of the mRNA vs. the level of translation revealed a population of mRNAs with marked induction of expression level, with no change at the level of translation, which were apparent mainly at the acute responses to all three stress conditions, as well as at the sustained response to heat shock (Fig. S1A). As we previously characterized widespread transcriptional readthrough in these conditions (Vilborg et al., 2017), we hypothesized that these may represent readthrough genes (also termed DoGs, Downstream of Genes-containing transcripts). However, mapping of DoGs using the DoGFinder tool (Wiesel et al., 2018), showed a very small overlap between DoG-generating genes and the aforementioned population. Instead, manual examination of individual genes from these populations showed that some of these genes are located downstream of readthrough regions, where the DoGs seem to continue into the downstream gene (Fig. 1A,B, Fig. S1B-E). Such a phenomenon has been documented before using analysis of nascent RNA following HSV-1 infection (Rutkowski et al., 2015), a condition known to induce transcriptional readthrough, and was termed “read-in” (Rutkowski et al., 2015). However, our data includes sequencing of mature, polyA-selected mRNAs. We therefore sought to characterize the read-in phenomenon more globally in this dataset. To that end, we set to identify read-in genes, which we defined as genes with substantial read coverage at the region 1kb upstream to the transcription start site (TSS) of the most upstream isoform of the gene, and were also located downstream of a readthrough event (a DoG, see Methods). This process identified overall 349 read-in genes, which were most abundant in the above three conditions, namely the acute responses to all three stresses, and in the sustained heat shock (Table S1). The sets of read-in genes showed significant overlap between the four conditions (Fig. 1C, 59 genes, p=1.44e-259), with higher overlap between the acute stress conditions (Fig. 1C, 73 genes, p=1.92e-192). Analysis of the distribution of distances between read-in genes starts and the 3’ ends of their upstream DoGs showed that, even though we did not require their proximity, in the vast majority of the cases (60-80%), they were indeed continuous with one another (Fig. S2). This indicated that most read-in genes were detectable as chimeric transcripts with their upstream DoG and readthrough gene.

**Figure 1:**
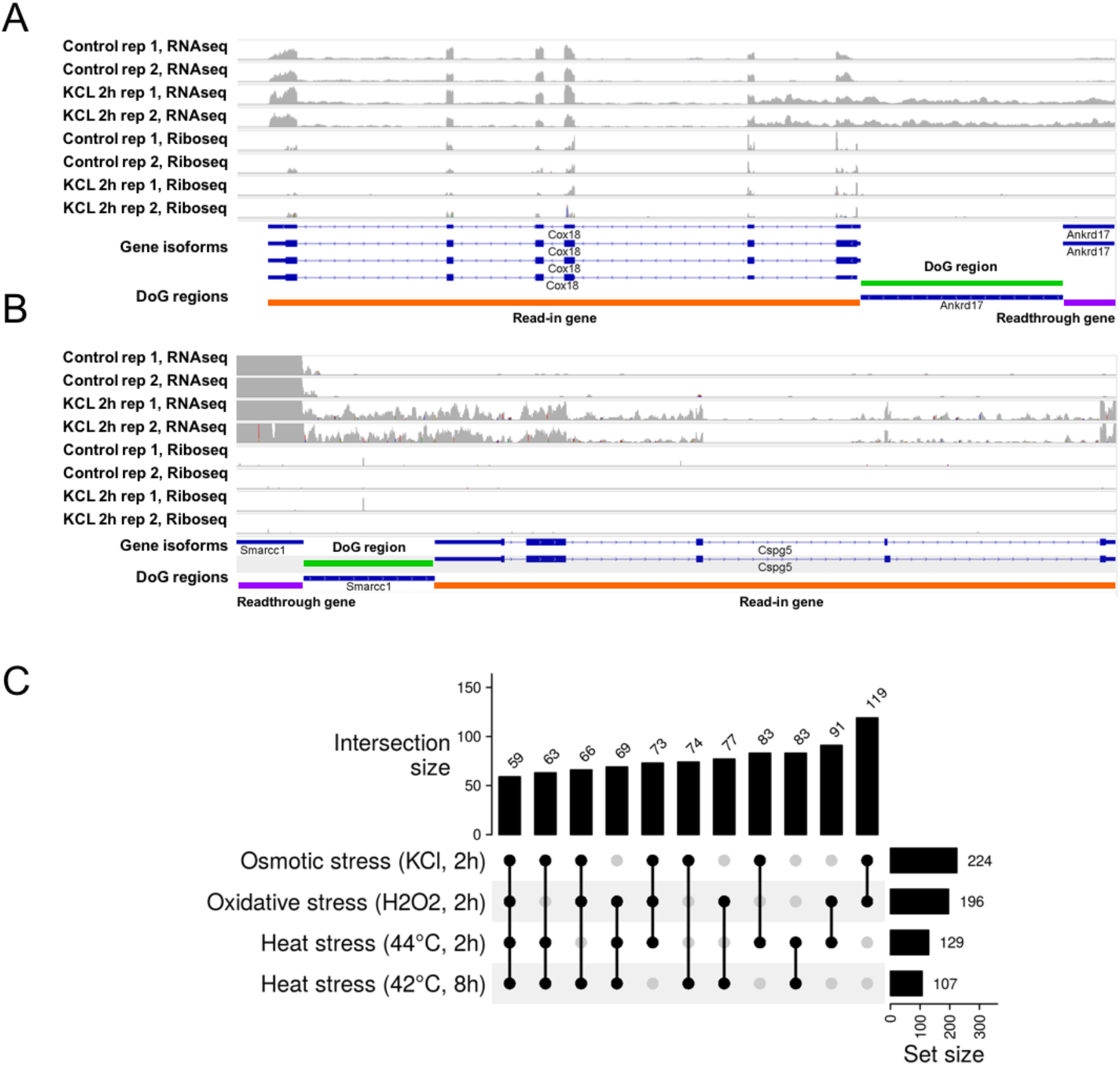
Pervasive readthrough into downstream neighboring genes in stress. (A-B) Read density plots (grey) for two examples of read-in genes, shown using IGV (Integrative Genomic Viewer 2.12.2(Robinson et al., 2011)), in control and osmotic stress (KCl, 2h) for expression (RNAseq) and translation (Riboseq). On the bottom, gene annotation tracks (in blue) are shown with the gene names, and a track representing the DoGs, i.e. the readthrough region, as generated using DoGFinder(Wiesel et al., 2018) is also presented (in blue). Regions of interest are highlighted with colors: the DoG region is highlighter in green, the readthrough gene in purple, and the read-in gene in orange. (C) Read-in gene sets are common between stresses. UpSet plot(Lex et al., 2014), visualizing the sizes of set intersections, of the overlap of the four stresses with the largest number of identified read-in genes, shows significant overlaps between them, with 59 read-in genes shared by all four (p=1.44e-259 of exact test of multiset intersection(Wang et al., 2015)).

### Read-in genes are exceptionally short and GC rich

Next, we sought to examine whether read-in genes demonstrate specific genomic characteristics. Examination of the distribution of distances to their upstream gene ends (on the same strand) showed that, as expected, read-in genes tend to be much closer to their upstream neighboring genes. Although shifted, the distributions of distances to the same-strand proximal gene for read-in and for DoGs without read-in overlapped (Fig. S3A). This suggested that read-in is not merely a consequence of proximity to a readthrough gene. We therefore wanted to identify additional properties that are characteristic of read-in genes. In order to control for potential confounding factors, we decided to compare the group of read-in genes to a control group of genes which were downstream of DoGs, and had a similar distribution of distances to their upstream readthrough gene ends, but were not classified as read-in genes in any of the conditions we tested. We termed this control group “non read-in genes”. Our first analysis showed that read-in genes tend to be substantially short (Fig. 2A); with a median length of about 6.1kb, which is less than a third of the median length of expressed genes (20.9kb). Non read-in genes showed an intermediate length, with a median of 14.5kb. Next, we asked in which gene regions this difference manifested, and, while exon lengths of read-in genes did not differ from the background of expressed genes (Fig. S3C), all other gene regions examined were significantly shorter (Fig. 2C, S3B,D,E). Most significantly, read-in genes had less introns (Fig. 2B), which were also much shorter (Fig. 2C), whereas non read-in genes had a similar number of introns to the entire population of expressed genes (Fig. 2B), with a median length of 0.7kb, which was much longer than that of read-in genes (0.3kbs), but shorter than the median population (1.2kb).

**Figure 2:**
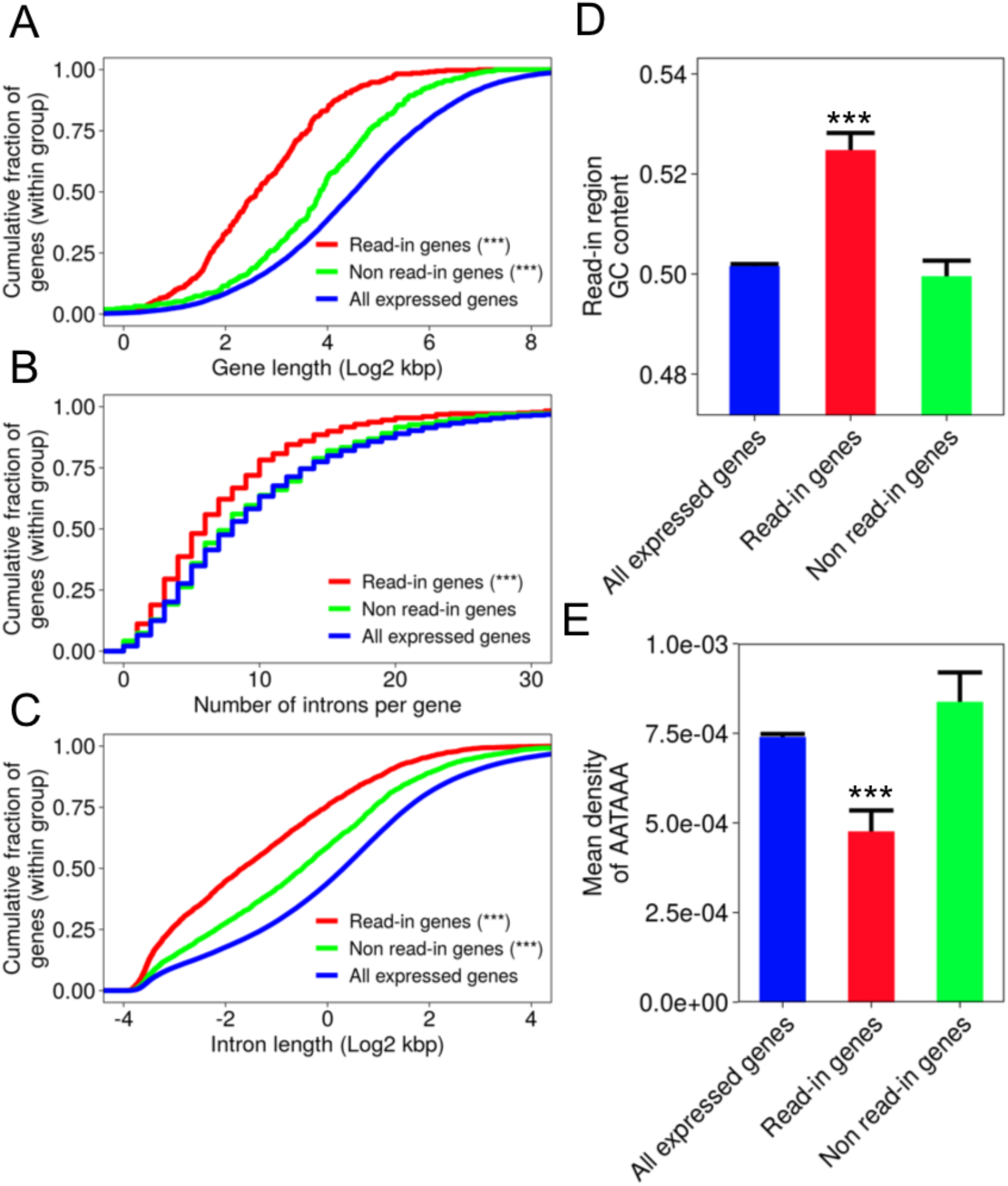
Read-in genes are significantly short, with fewer shorter introns, and GC rich. (A-C) Distribution of feature lengths (log2 kbp) of read-in (red), non-read-in (green) and all expressed genes (blue) presented as cumulative distribution function (CDF) plots. p-values were calculated using Wilcoxon rank sum test, between either read-in or non read-in distributions vs. all expressed genes, *** p<0.001, p indicated when significant (p<0.05). Shown are the length of the entire gene (A), p(read-in vs. expressed) = 1.9e-78, p(non read-in vs. expressed)=2.27e-11, number of introns per gene (B), p(read-in vs. expressed) = 2.8e-10, p(non read-in vs. expressed)=6.38e-01, and introns lengths distributions (C) p(read-in vs. expressed)<2.23e-308, p(non read-in vs. expressed)=4.24e-05). (D) GC content was significantly higher in the 1kb upstream regions, p-values were calculated using Wilcoxon rank sum test, *** p<0.001, p(read-in vs. expressed)=2.86e-11, p(non read-in vs. expressed)=8.32e-01. (E) PolyA signals are significantly depleted in read-in regions (1kb upstream regions), compared to the corresponding regions upstream to all expressed genes. p(read-in vs. expressed)=2.8e-09, p(non read-in vs. expressed)=6.71e-01. Two additional variants of the polyA signal show similar trends, Fig. S3H,I. p-values were calculated using Wilcoxon rank sum test, *** p<0.001.

Next, we focused on the “read-in region”, i.e., the 1kb region upstream to read-in genes, and found that this region is substantially GC rich, compared to both the background of all expressed genes as well as non read-in genes (Fig. 2D). Read-in genes also had GC rich introns (Fig. S3F) and were overall significantly GC richer than expressed genes (Fig. S3G).

Previous studies have shown that DoG regions tend to be depleted for polyA signals (Rutkowski et al., 2015; Vilborg et al., 2015; Vilborg et al., 2017), reasoning that multiple polyA signals could assist in efficient termination in times when the termination is partially impaired. We thus analyzed the polyA signal content in the read-in regions, and found that, here too, polyA signals were significantly depleted compared to the upstream regions of all expressed genes, as well as those of non read-in genes (Fig. 2E, S3H,I).

### Genes with a high degree of read-in are translationally inhibited in stress conditions

Having observed that several examples of read-in genes show low levels of translation in stress (Fig. 1B, Fig. S1B-E), we asked if this trend could be generalized for the entire group of read-in genes. Our analyses show that, following acute osmotic stress, read-in genes showed a significant induction at the mRNA level, however their translation was unchanged (Fig. S4A). In acute heat shock and oxidative stress, read-in genes were induced at the level of expression, while at the level of translation, a significant but milder effect, on average, was observed (Fig. S4B,C). Next, we asked whether read-in is associated with a translational shutoff during stress. A comparison between mRNA expression and translation levels showed that only a subset of the read-in genes is translationally inhibited (Fig. 3A, S4D). Manual examination of several examples indicated that, in some cases, the expression of the upstream DoG seemed to be similar to that of the read-in gene (as in Fig. 1B, S1B,C), while in other cases, the expression of the read-in gene was clearly higher than that of its upstream DoG (as in Fig. 1A, S1E). We therefore sought to quantify this effect for all read-in genes, reasoning that read-in genes transcripts with similar levels of expression as their upstream DoGs are probably mostly expressed due to readthrough transcription, while those with a higher expression than their upstream DoG represent a mix of readthrough transcription and independent expression. We therefore calculated a metric termed “read-in estimation”, which is simply the read density in the read-in region (1kb upstream to the TSS) divided by the read density over the entire mRNA (see Methods). Stratifying read-in genes into three equal-sized groups according to their read-in estimation values showed that the subset of genes with high read-in estimation values, meaning that most of their expression could be attributed to readthrough transcription, had the lowest levels of translation in all conditions (Fig. 3A-B, S4D-E). Furthermore, the lower the read-in estimation was, the higher the translation levels were in all conditions (Fig. 3A-B, S4D-E). To understand whether these translation levels could merely reflect the expected translation given the levels of the mRNA, we calculated the ratios between translation and expression levels (often termed “Translation Efficiency” or “TE”) and compared the mean TE values of high, medium, or low read-in estimation genes to the mean TE values of randomly sampled groups of mRNAs with expression levels that were matched to those within each read-in gene group (see Methods). We found that, while for read-in genes with low read-in estimation values, translation levels were no different than expected given their corresponding mRNA levels, read-in genes with high read-in estimation values were substantially less translated than expected given their expression levels in all conditions (Fig. 3C, S4F). Genes with intermediate levels of read-in showed significantly lower translation-to-expression ratios than expected in the three acute stress conditions, and in sustained heat shock, but not in other sustained stresses or control conditions (Fig. 3C, S4F). Therefore, read-in genes with high read-in estimation, i.e. for which expression is mainly due to readthrough, are indeed translationally repressed.

**Figure 3:**
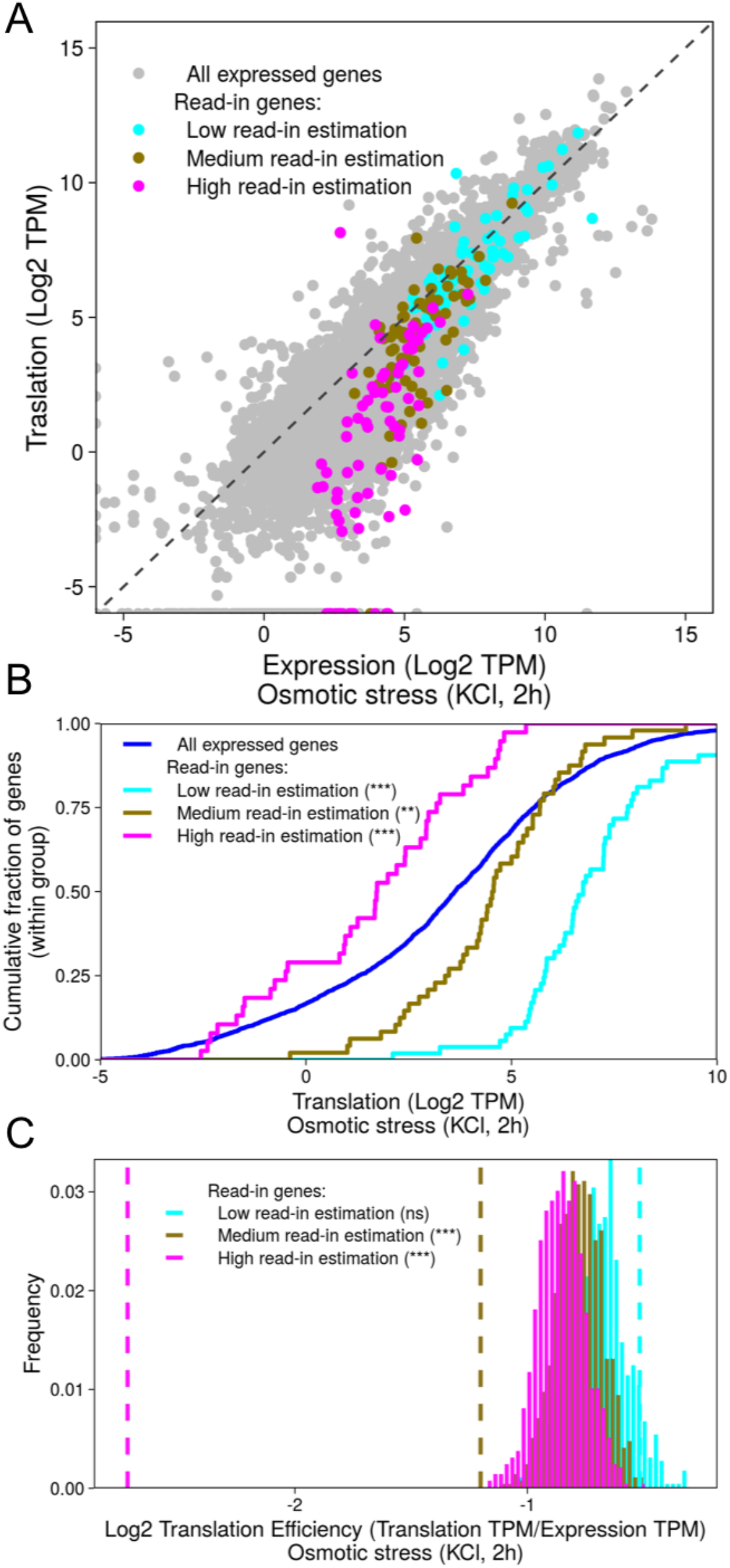
Read-in genes tend to be lowly translated. (A) Scatter plot of Log2 TPM values of expression and translation of read-in genes during osmotic stress (2h). Read-in genes were stratified by their read-in estimation levels (cyan, brown and magenta) corresponding to 33% quantiles of read-in estimation values (see Methods). All expressed genes are shown in grey. Similar trends are also shown in other conditions (Fig. S4D). (B) CDF plot of the TPM translation values (in log2) of read-in estimation groups in Osmotic stress, 2h shows an inverse correlation between the degree of read-in and the level of gene translation. p-values were calculated using Wilcoxon rank sum test, *** p<0.001, **p<0.01, (see Table S2 for exact p-values). Similar trends are also shown in other conditions (Fig. S4E). (C) Random sampling analysis of translation efficiency (translation normalized to expression), where randomly sampled groups were matched for levels of expression with their corresponding read-in estimation group (high, medium and low are color-coded, see Methods) in Osmotic stress (2h). dashed lines represent the mean translation efficiency value for the read-in group. The analysis demonstrates a significantly lower translation efficiency for high and medium read-in estimation genes (*** - p<0.001 for both, see Table S2 for exact p-values) in light of the expected translation levels given their expression levels (solid colored distributions). Similar trends are also shown in other conditions (Fig. S4F).

### Read-in genes show marked intron retention

Looking at several individual examples of read-in genes, we noticed that they show high levels of intron retention of their first intron (Fig. 1A,B, S1B-E), and occasionally also for other introns (Fig. 1B,S1C). We therefore asked how prevalent intron retention is in read-in genes, in a systematic manner. We first calculated the PSI value (Percent Spliced In) for each intron in the genome, as a metric to evaluate the extent of intron retention (see Methods). Indeed, we found that, overall, read-in genes show marked intron retention compared to other genes (Fig. 4A, S5A), in all conditions. Furthermore, intron retention was not restricted to first introns, although the highest levels of intron retention were found in first introns (Fig. S5B). Interestingly, the higher the read-in estimation levels were, the higher the degree of intron retention that was observed (Fig. 4A, S5A). Since intron retention is known to lead, in many cases, to nuclear retention (Monteuuis et al., 2019; Shalgi et al., 2014), this could explain why genes with high read-in estimation levels are lowly translated. Examination of all three factors together; read-in estimation, intron retention, and translation levels, showed that, indeed, the higher the degree of intron retention was, the lower the translation levels were, and the higher the read-in estimation was for read-in genes (Fig. 4B, S5C).

**Figure 4:**
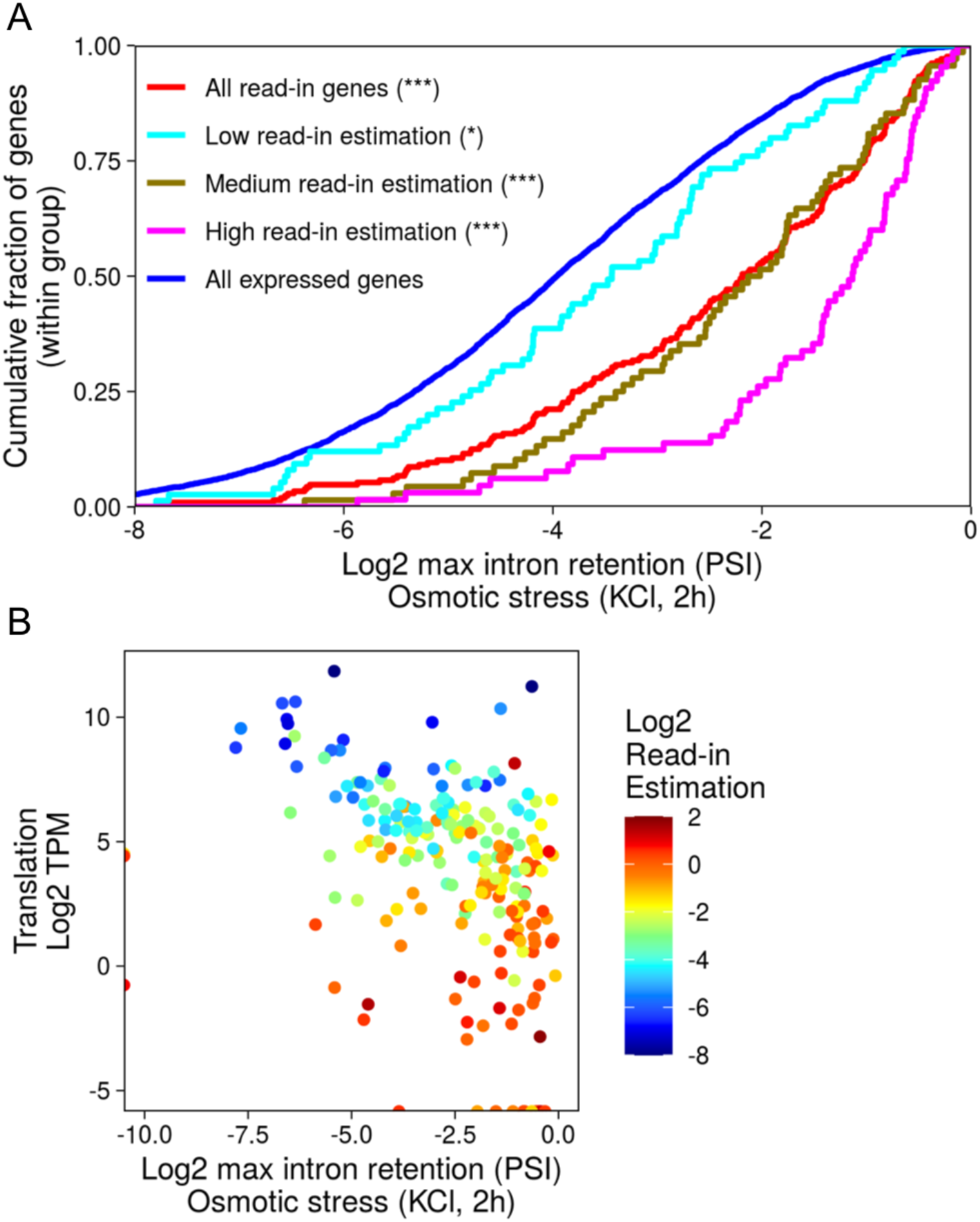
Read-in genes show marked intron retention. (A) CDF plot of maximal intron retention value per gene (PSI, in log2, see Methods) in osmotic stress (2h), shown for all read-in genes (red) or split into three different read-in estimation groups (cyan, brown and magenta), compared to a background set of all expressed genes (blue), demonstrate significantly higher degrees of intron retention for all groups, that increase with the degree of read-in estimation. Wilcoxon rank-sum test p-value of each group vs. all expressed genes, *** p<0.001, *p<0.05 see Table S2 for exact p-values. Similar trends are also shown in other conditions (Fig. S5A). (B) Scatter plots of translation levels (log2 TPM, y-axis) vs. maximal intron retention (log2 PSI value, x-axis) for read-in genes in osmotic stress (2h) exhibit a significant negative correlation (R = -0.482, p(R) = 5.47e-05). The color axis shows the read-in estimation values (in log2), further demonstrating that the more the intron is retained the higher the read-in estimation tends to be (R = 0.493, p(R) = 0.000502). Similar trends are also shown in other conditions (Fig. S5C).

### Intronic features in read-in genes make them prone to intron retention

We found that read-in genes have particularly short introns, which were significantly GC rich (Fig. 2C, S3F). It was previously shown that these two properties, i.e. short intronic length and high GC content, are associated with an increased tendency for intron retention (Monteuuis et al., 2019). We analyzed the entire set of introns, and indeed observed that these tendencies are largely recapitulated in our data for the entire set of introns (Fig. S6A). We therefore asked whether these properties alone could explain the marked degree of intron retention that we observed in read-in genes. To answer this, we performed a random sampling test, where sets of introns with the same length and GC content distribution as those in read-in genes were randomly sampled, and the mean degree of intron retention was calculated for each random set (see Methods). This would allow us to specifically control for the two intronic properties known to be associated with intron retention. Surprisingly, this analysis showed that these two features could not explain the high degree of intron retention observed in read-in genes, which was significantly higher than expected in all conditions (Fig. 5A, S6B), and in particular in the acute stress conditions.

**Figure 5:**
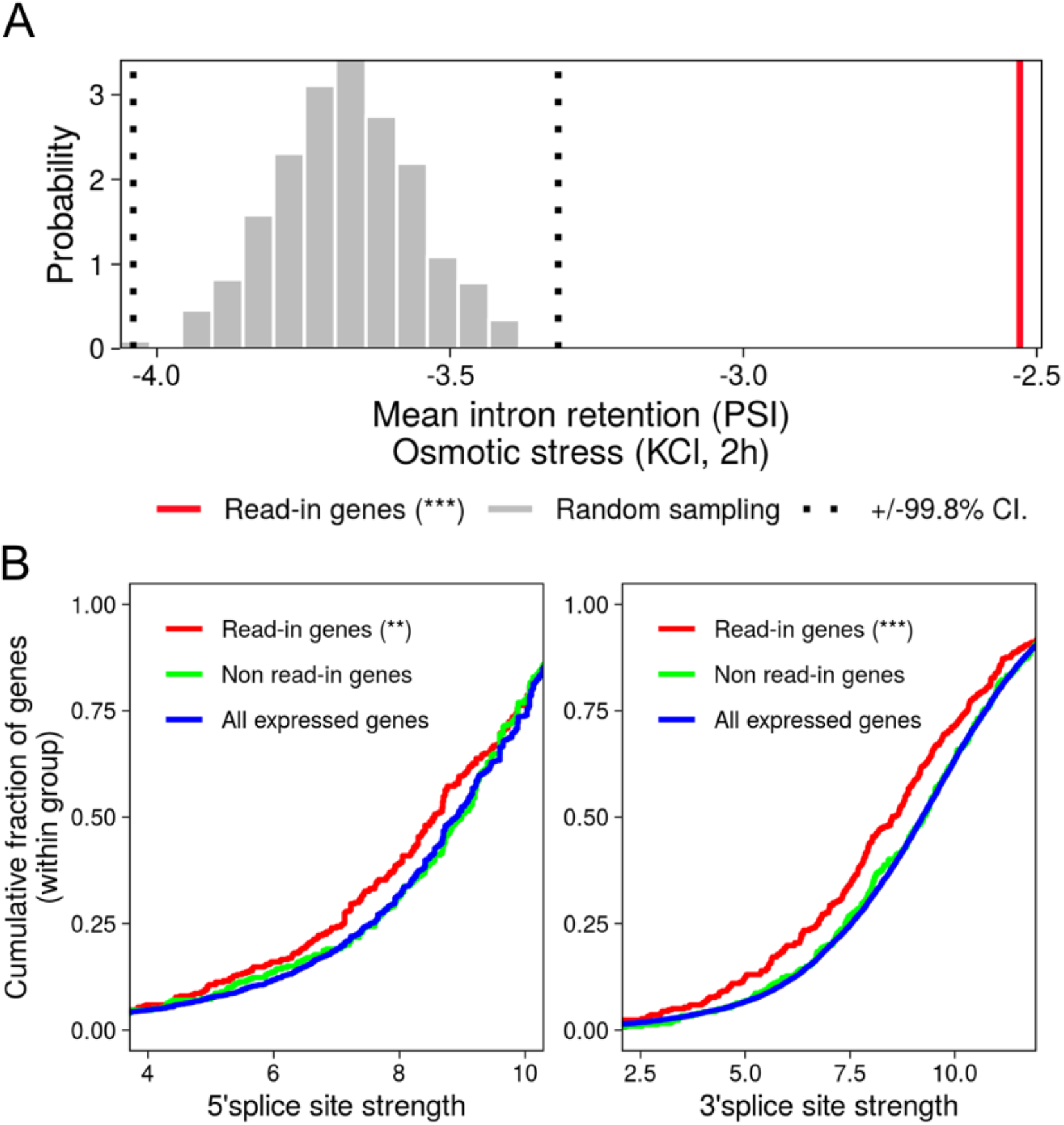
Read-in genes intron retention levels are much higher than expected given their genomic characteristics. (A) A thousand randomly sampled comparison groups, matched for GC content and intron lengths distributions as in read-in genes identified in osmotic stress (2h) were generated (see Methods). For each randomly sampled comparison group, the mean value of the maximum intron retention (log2 PSI) per gene was calculated and plotted as a histogram (grey). Confidence intervals (+/-99.8%, corresponding to 3*std) are presented as dashed black lines. The mean value of the maximum intron retention (log2 PSI) of read-in genes (red) is significantly higher than the distribution of the mean intron retentions, even when controlling for GC content and intron lengths (*** p < 0.001). Similar trends are also shown in other conditions (Fig. S6B). (B) CDF plots of 5’ (left) and 3’ (right) splice site strengths, calculated as MaxEnt scores (Yeo and Burge, 2004) (see Methods), show significantly weaker splice site strengths distribution for first introns in read-in genes (red) compared to all expressed genes (blue), while non read-in genes splice site strengths (green) are similar to those of all expressed genes. p-values were calculated using Wilcoxon rank-sum test, *** p<0.001, ** p<0.01. For 5’ splice sites p(read-in vs. all expressed)=0.009, p(non read-in vs. all expressed)=0.85, for 3’ splice sites p(read-in vs. all expressed)=3.46e-06, p(non read-in vs. all expressed)= 0.72.

We thus turned to examine another property which may partly explain the extreme intron retention that we observed in read-in genes, namely splice site strength (Yeo and Burge, 2004) (see Methods). We quantified splice site strength for all first introns using MaxEnt (Yeo and Burge, 2004). Our analysis showed that first introns in read-in genes have significantly weaker splice sites, both 5’ as well as 3’ splice sites, compared to the entire distribution of first introns, while first introns in non read-in genes were no different than the entire population (Fig. 5B). Thus, first introns in read-in genes are not only short and GC rich, but also have particularly weak 5’ and 3’ splice sites.

## Discussion

Under stress conditions, reduced termination efficiency generates widespread readthrough transcription (Rosa-Mercado and Steitz, 2021). Here we showed that this phenomenon leads to pervasive read-in into downstream neighboring genes, which tend to have short introns and high GC content, and present marked intron retention.

Previous studies found that housekeeping genes are generally short (Eisenberg and Levanon, 2003). However, read-in genes did not significantly overlap with the set of housekeeping genes (hypergeometric p>0.999, housekeeping genes taken from (Eisenberg and Levanon, 2003)), and pathway analysis showed no significant enrichment with any particular functional category. Interestingly, early studies highlighted a correlation between GC-content and gene-dense genomic regions (Bernardi, 2000). Nevertheless, when we compared the set of read-in genes to non read-in genes, which have a similar gene proximity distribution to that of read-in genes, we observed that upstream regions of non read-in genes have the same GC content as that of the expressed genes background (Fig. 2D). When looking at the gene itself, the GC content of non read-in genes was higher than that of the expressed genes background, but still significantly lower than the GC content of read-in genes (Fig. S3G). Thus, read-in genes seem to be GC richer than expected even given their tendency to be proximal to their upstream genes.

Our data showed that read-in is associated with intron retention (Figs. 4,5). This process may be combined with splicing inhibition under stress, as was shown to occur in heat shock (Shalgi et al., 2014), to increase intron retention even further. Interestingly, in budding yeast, intron retention, readthrough and read-in transcription were recently shown to occur in a knock-out strain of Nab2, an essential poly(A) binding protein involved in splicing and nuclear export (Alpert et al., 2020). These further highlight potential mechanistic links between read-in and splicing defects, as we present here, during naturally-occurring stress-inducing, readthrough promoting conditions.

Intron retention was shown to be conserved in evolution, and to be correlated with proteome complexity (Schmitz et al., 2017). Furthermore, intron retention in mRNAs may lead to a variety of fates, most of them resulting in translation inhibition (Monteuuis et al., 2019). In rare cases, intron-containing mRNAs might be exported from the nucleus, where they are subjected to NMD-mediated decay (Lykke-Andersen and Jensen, 2015). In a few instances, retained-intron mRNA (termed Detained-introns) were shown to be spliced in response to specific stimulations (Boutz et al., 2015; Tan et al., 2020). However, the most common fate of intron-retained mRNAs is nuclear retention (Monteuuis et al., 2019), where they may subsequently be degraded in an NMD-independent manner (Yap et al., 2012). Here, with data representing a steady-state snapshot of polyA-selected, mature, whole-cell mRNAs, it seems that read-in transcripts were not degraded post maturation, but rather were stable enough to allow detection as mature, polyadenylated mRNAs by RNA-seq. It is possible that read-in has co-evolved with genomic properties that favor extremely high degrees of intron retention, thereby aiding the nuclear retention of read-in gene transcripts during readthrough-promoting stress conditions. Our findings suggest that read-in, and perhaps also its upstream readthrough, are not merely a by-product of stress-induced failure in polyadenylation and transcription termination, since the properties of read-in genes may facilitate the quality control for read-in transcripts, ultimately allowing stress-induced readthrough transcription to persist, while preventing the nuclear export of read-in transcripts, and precluding their unwarranted translation.

## Methods

### Datasets

Transcriptomics, using polyA selected RNA sequencing, and translatome mapping, using Ribosome footprint profiling, were conducted in duplicates, in two separate experiments, each having its own respective control. NIH3T3 mouse fibroblast cells were exposed to Heat shock (44°C for 2 hours or 42°C for 8 hours), or remained at 37°C as control. Additionally, NIH3T3 were exposed to Oxidative stress (0.2mM H2O2 for 2 or 7 hours) and Osmotic stress (80mM KCL for 2 or 7 hours). Libraries were made using the NEB E7760 kit, and ribosome footprint profiling was performed as in Sabath et al.(Sabath et al., 2020).

### Gene annotation

NCBI Refseq mm10 and UCSC RefSeq mm10 tables were downloaded from the UCSC genome browser website, using UCSC Table Browser data retrieval tool. To facilitate discovery of DoGs and read-in regions, NCBI Refseq annotation of mm10 mouse genome was combined with the UCSC Refseq annotation, by adding the isoforms unique to UCSC Refseq to the NCBI Refseq annotation. Annotations of non-translated short RNAs, such as snoRNAs, miRNAs and pseudogenes, were filtered out.

### RNAseq and Riboseq analysis – initial read mapping and transcript quantification

For RNAseq analysis, preliminary filtering of sequences was performed. The raw reads were mapped to sequences of rRNAs using STAR(Dobin et al., 2013) aligner, with all mapped reads discarded. Following the filtering, the remaining raw reads were mapped to mm10 mouse genome, using the STAR aligner with the following parameters: --outFilterMatchNminOverLread 0.6. Transcriptome sequences were taken from the mm10 mouse genome annotation described above.

For Riboseq, preliminary filtering of non-mRNA footprint sequences was first performed. Raw reads post adapter trimming were mapped to sequences of rRNAs, tRNAs and snRNAs, using STAR aligner, with all mapped reads discarded. Following the filtering, the remaining reads were mapped to the coding sequences of the mm10 mouse genome annotation described above, which were clipped by 30 nucleotides from start and end of each CDS, since there is a positional bias of the ribosome in those areas. The mapping was done using the Bowtie 2 aligner(Langmead et al., 2009) with the following parameters: -- outFilterMatchNminOverLread 0.6. After alignment of RNAseq and Riboseq samples, quantification was performed using the RSEM software package(Li and Dewey, 2011). RSEM returns a raw read count and a normalized TPM (Transcripts Per Million) value for every gene and every isoform in the annotation. Expressed genes were defined as genes that had an RNAseq TPM value larger than 4 in at least one sample.

### Differential expression analysis of RNAseq and Riboseq

Differential expression analysis of transcriptome and the translatome data was performed on read counts data from the output of RSEM, using DESeq2 R package(Love et al., 2014), with the following parameters: cooksCutoff=FALSE, independentFiltering=TRUE, lfcShrink=TRUE. The comparison was performed between each stress condition samples and their corresponding controls. Log2 Fold Change (LFC) values returned by DESeq2 package were used in further analyses.

### Identification of DoGs

Sorted bam files produced by RNAseq STAR genomic alignment were used as input for the DoGFinder tool(Wiesel et al., 2018) in order to identify DoG regions. The annotation described above was used. Get_DoGs function was run with the following parameters: -S -minDoGLen 100 -mode W –minDoGCov 0.6. DoGs RPKM were calculated using the DoGFinder tool. DoGs were filtered to express RPKM value of at least one and a total number of mapped reads of at least five.

### Read-in genes

For each DoG-producing gene, a downstream read-in region was defined as a 1kbp 5’ flank region of the most upstream isoform of the proximal downstream gene on the same strand. In cases where the distance between genes was less than 1kpb, the entire region between the two genes was used. Coverage and read counts of the read-in regions were calculated using Bedtools(Quinlan and Hall, 2010). Read-in genes were defined as genes with the read-in region RPKM value of at least one, coverage of at least 0.6, and a TPM value of the downstream gene of at least four. A set of all read-in genes was defined as all genes that were defined as read-in in at least one sample, resulting in a total of 349 read-in genes. To define read-ins occurring under a particular stress, a union of the read-ins of the two replicate samples of that stress was used.

### Non-read-in genes

A set of non-read-in genes was defined as a control group, by taking a set of all genes that were located downstream of DoG-containing genes on the same strand, but were not defined as read-in in any of the samples. In addition, to further control for confounding factors related to gene proximity, these genes were filtered by their distance from the end of the DoG containing genes, such as that the distance was lower than the 95% percentile of the distance distribution between read-in genes and their upstream readthrough genes (a distance less than 11644 bp), resulting in a total of 321 non-read-in genes.

### Feature analysis of gene sequences

For the analysis of isoform length, coding sequence (CDS) length, exon length, intron length, 5’ and 3’UTR lengths, number of introns and GC content of introns, the isoform most expressed in the RNAseq control samples, as calculated by RSEM, was selected for each gene.

For the analysis of read-in regions GC content and polyA signals, the read-in region was used, defined as a 1kbp 5’ flank region of the most upstream isoform of the gene. In cases where the distance between genes was less than 1kpb, the entire region between the two genes was used.

### Read-in estimation values

For each gene downstream of a DoG-containing gene, and for each sample, read-in estimation value was defined as read density of the read-in region divided by the read density of the most highly expressed isoform of the gene in the sample. Read-in estimation value for a given stress was defined as a mean of the values of the two replicate samples.

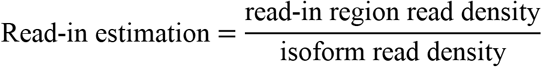

### Intron retention

For each intron and for each sample, number of exon junction reads and intron reads was extracted using Bedtools software(Quinlan and Hall, 2010) and read density was calculated. PSI (percent spliced in) was calculated using the following formula:

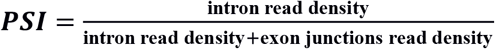

Introns that had exon junction read density of zero were discarded. For each gene in each sample, the most highly expressed isoform was selected, and the intron with the highest PSI value was selected to represent maximum intron retention for this gene. For analysis of first intron retention, the first intron of the most highly expressed isoform was selected. Intron retention for a given sample was defined as a mean of the PSI values in the two replicate samples.

### Splice site strengths analysis

For splice site strength analysis, the most highly expressed isoform for each gene was chosen, and a custom R script was used to generate fasta files of splice site sequences of the first intron of the isoform. MaxEnt software was used to calculate the 5’ and 3’ splice site strength scores(Yeo and Burge, 2004).

### Analysis of correlation between read-in intensity and translation efficiency

To assess the changes in the translation efficiency within read-in genes compared to expressed genes, translation efficiency (TE, defined as translation TPM values divided by expression TPM values) was calculated for each gene in each sample. TE of a gene for a given stress was defined as a mean of the TE values in the two replicate samples. For each of the three read-in estimation groups (low, medium and high), mean log2 TE was calculated, and groups of 1000 randomized samples of TE values were taken from the set of expressed genes, controlling for matching range of TPM values of the read-in genes within the specific group. P-values were calculated by comparing the distributions of the mean log2 TE values of the randomized samples to the mean values of each of the read-in estimation groups.

### Controlled analysis of intron retention within read-in genes

Read-in genes were divided to 16 groups according to a combination of 4 quantiles of intron lengths and 4 quantiles of intronic GC content. Then, 1000 randomized comparison groups with the same GC content and intron lengths distributions as read-in genes were picked from the set of all expressed genes without read-in genes. For each comparison group, the mean value of the maximum intron PSI for each gene was calculated.

## Data availability

All RNA-seq and ribosome footprint profiling data were deposited in GEO, accession number GSE197536. Ribosome footprint profiling data for heat shock was taken from GSE32060.

## Acknowledgements

We thank members of the Shalgi lab for fruitful discussions throughout the project. We thank Flonia Levy-Adam and Naseeb Saida for critical reading of the manuscript.

